## Supplementary_Figures_Video_Captions for "Actin nano-architecture of phagocytic podosomes"

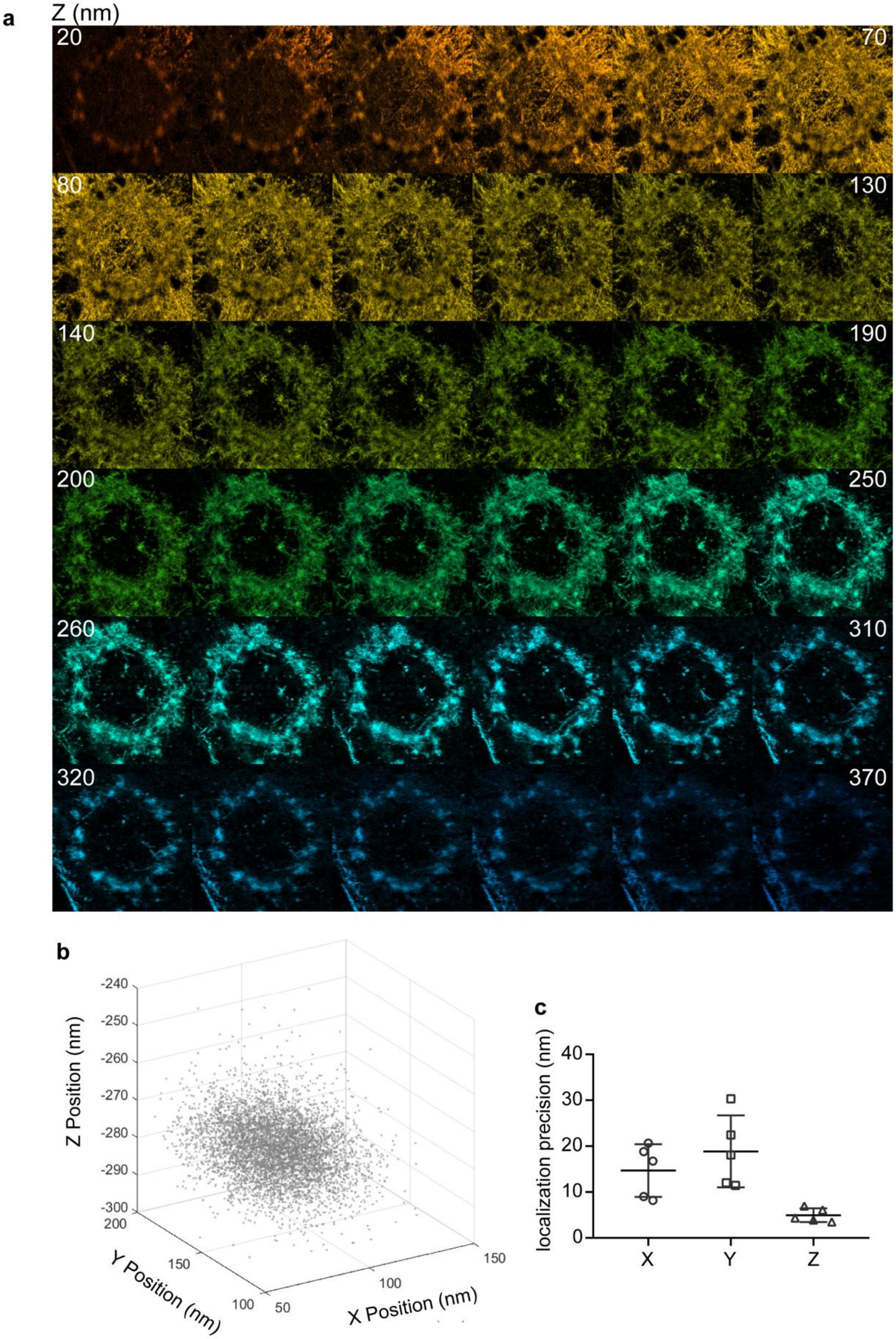
a
Z (nm)
20
70
80
130
140
190
200
250
260
310
320
370
b
Z Position (nm)
-240
-250
-260
-270
-280
-290
-300
Y Position (nm)
200
150
100
X Position (nm)
50
100
150
c
localization precision (nm)
40
30
20
10
0
X
Y
Z

**Supplementary Figure 1. Additional information for iPALM imaging. a)** Additional images from the Z-stack of the frustrated phagocytosis site shown in Fig. 2a,c. Snapshots are shown at 10 nm height intervals. **b, c)** Localization precision of the iPALM system. Localization of one gold nanorod fiducial along time (randomly selected 6000 frames) is shown in **b**. Localization precision was estimated from the histogram of nanorod positions and inferred by the standard deviation (σ = FWHM/2.3) of the distribution for each axis (shown in **c**).

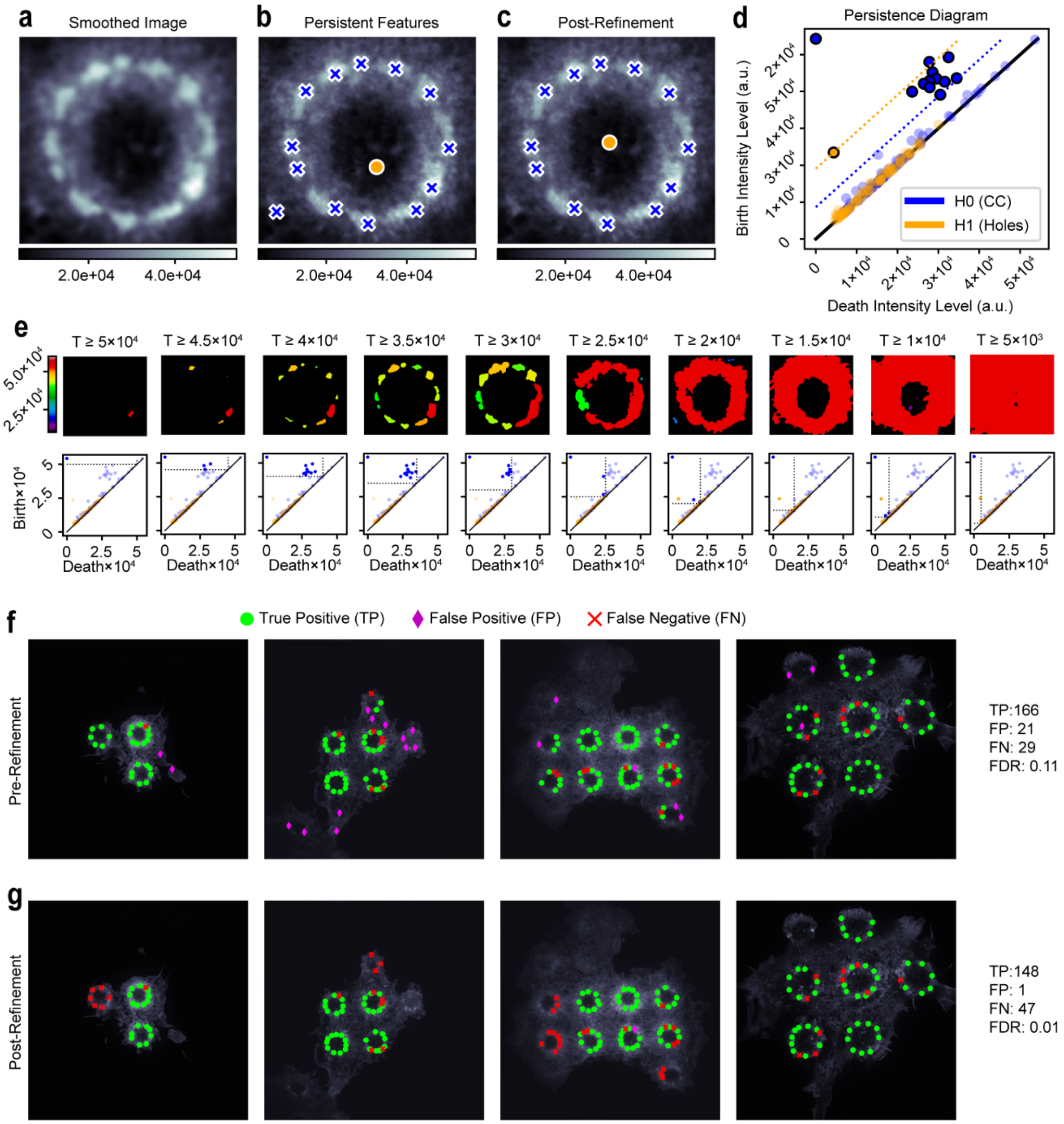


**Supplementary Figure 2. Persistent homology methods for identifying podosomes and site locations. a)** Smoothed image of a single phagocytosis site. An insignificant amount of noise is added for uniqueness. **b)** Locations of significantly persistent features from the pipeline. **c)** Final locations of podosomes and the phagocytosis site centers after post-processing that excludes podosomes far away from phagocytosis sites and adjusts the locations of phagocytosis site centers. **d)** Persistence diagram based on image from a. Pixels equal to the birth levels of significantly persistent features (dots to the upper left of the persistence threshold, dotted lines) in h0 and h1 (connected components and enclosed holes respectively) are shown in b. **e)** Demonstration of level-set filtration. Visible connected components (top, colored by maximum value) and the persistence diagram (bottom, visible features correspond to top left quadrant) are shown as the level drops. **f,g)** Podosomes were manually identified if empirically determined to be associated with a well-defined phagocytosis site with at least 3 podosomes. True positives were located both manually and by the identification pipeline, false positives were located by the pipeline but not manually, and false negatives were located manually but not by the pipeline. Results are shown both before (**f**) and after (**g**) the refinement step is performed.

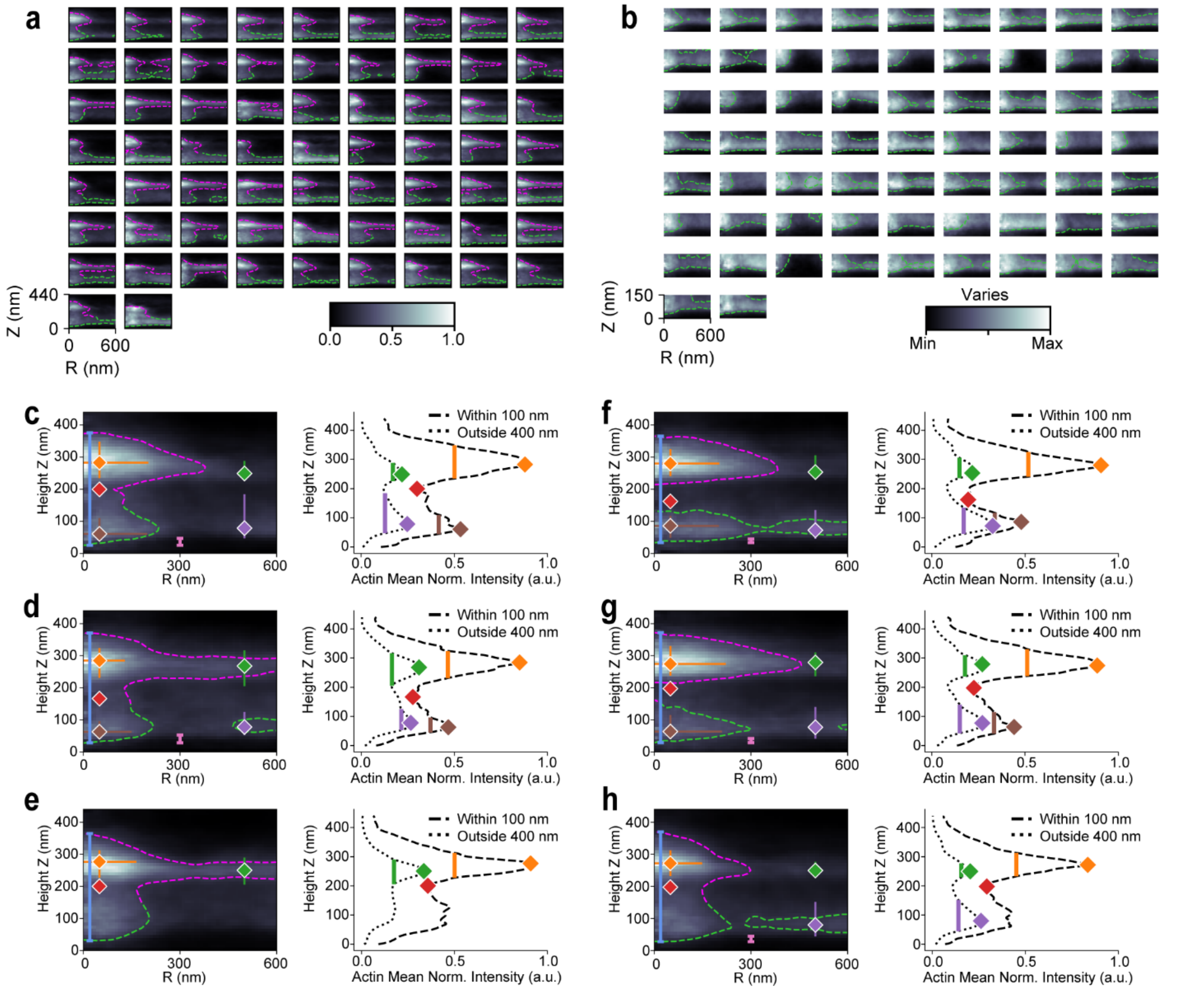


**Supplementary Figure 3. Heatmaps of quantified podosomes. a)** All individual radial averaging heatmaps from iPALM data, except those shown in Fig. 4f,g (n = 66 podosomes). Magenta contour based on mean actin intensity within a radius of 350 nm. **b)** All individual zoomed (Z 0 – 150 nm) radial averaging heatmaps from **a**. Each zoomed heatmap corresponds to the heatmap in the same row and column in **a**. **c-h)** Additional examples of features extracted from individual radial averaging heatmaps. These examples correspond to Fig. 4d,e.

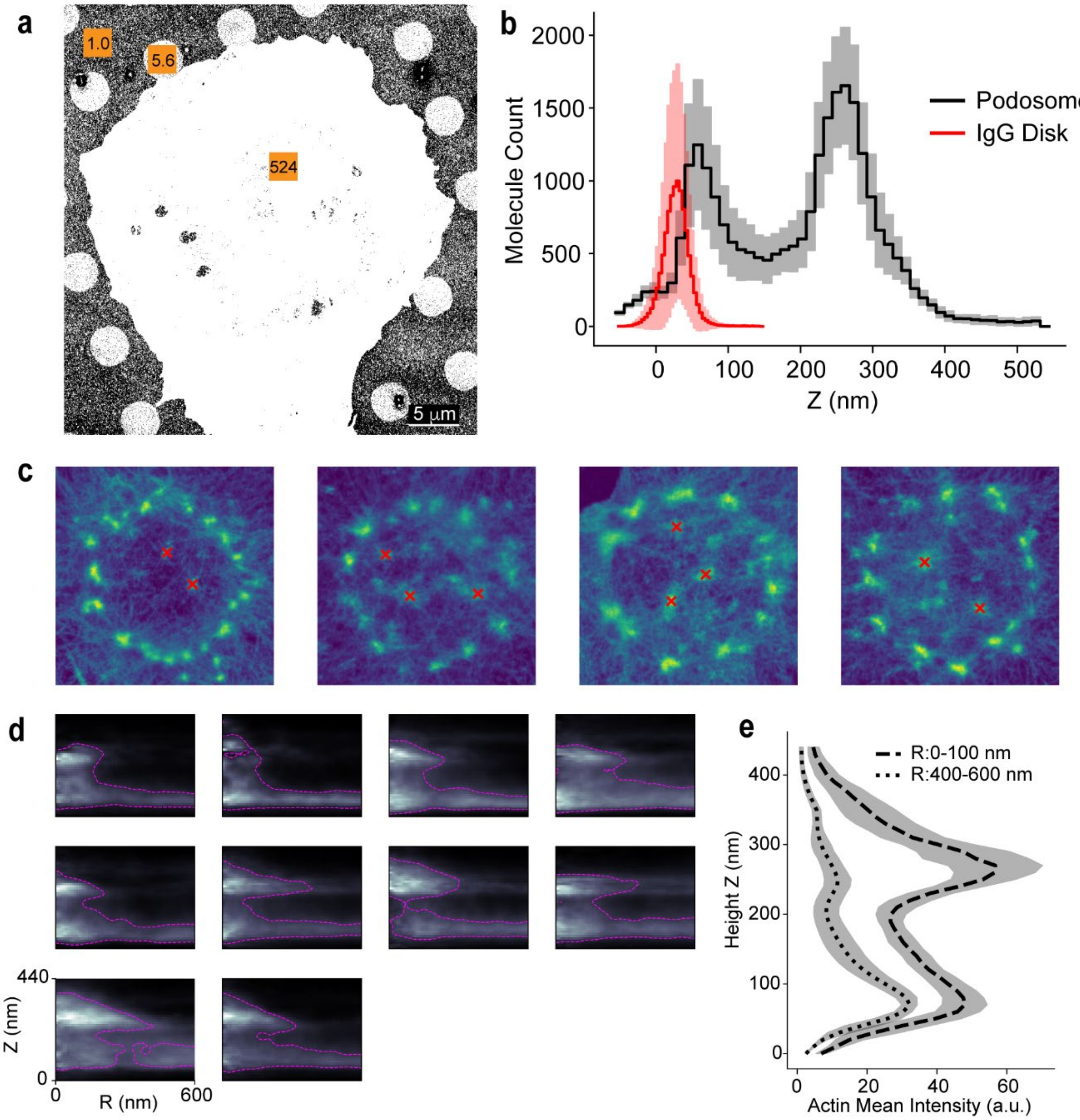


**Supplementary Figure 4. Control experiments to demonstrate that IgG topology did not influence podosome morphology. a)** Off-target binding of Phalloidin Alexa 647 to IgG was used for iPALM imaging of IgG disks. In the orange boxes, the normalized molecule count per $\mu m^2$ is shown for the background, within the cell, and within the IgG disks. **b)** The IgG disk (red, n = 11) is low and narrow compared to the neck and other major features of podosomes (n = 5), indicating that the bi-lobed structure of the podosome is not an artefact of formation around the edge of the IgG. **c)** iPALM imaging of podosomes that formed over IgG disks ($n_{sites}$ = 4, $n_{pod}$ = 10), rather than at edges. **d**) Radial averaging heatmaps of the podosomes in **c**. Magenta contour is drawn at 70% of the maximum intensity. **e)** Within 100 nm and from 400-600 nm, mean actin intensity for the heatmaps in **d**.

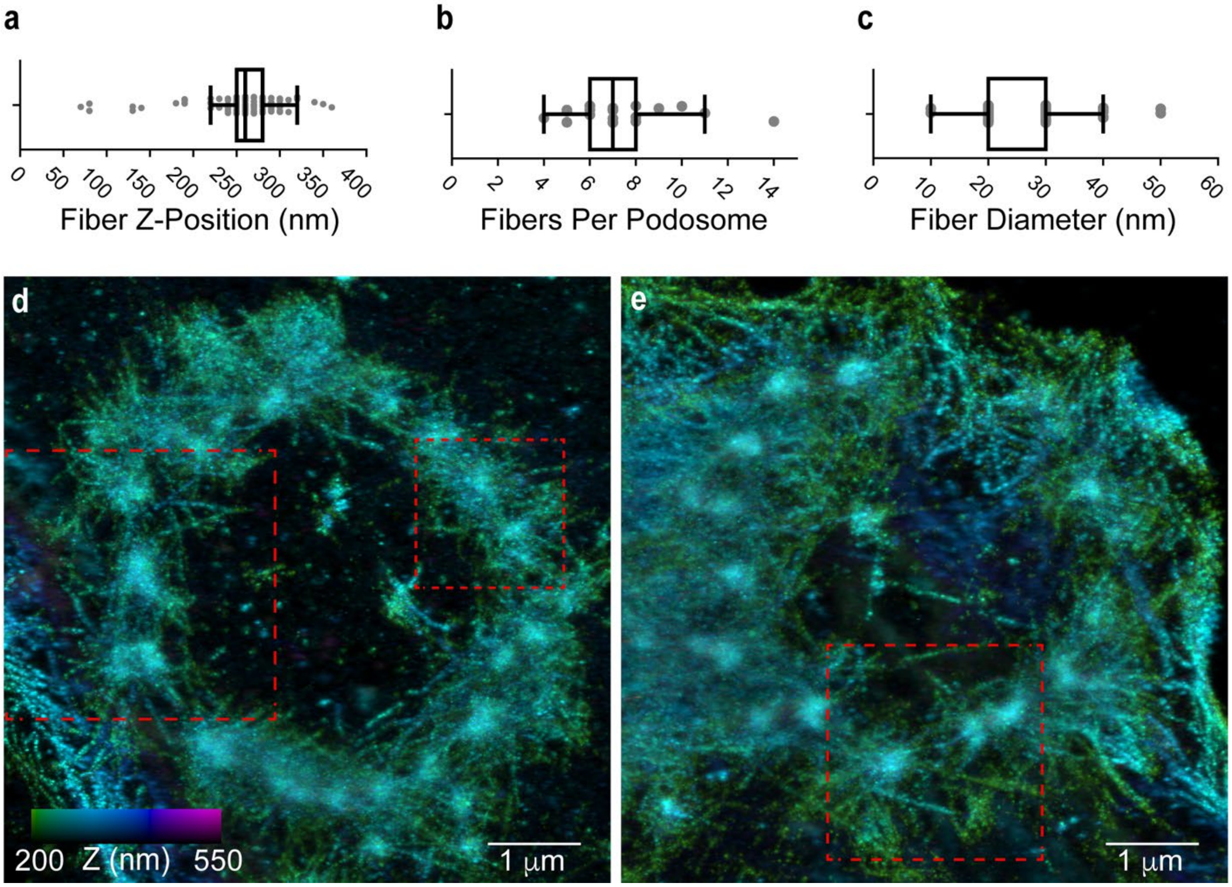


**Supplementary Figure 5. Further characterization of radial filaments. a-c)** Average Z-position for radial filaments, the number of filaments per podosome, and filament diameter. Measurements were extracted from Imaris analysis (n = 146 filaments across 20 podosomes). **d,e)** From two rings of podosomes, examples of color coded Z-projections. Color bar indicates z distance from 200 to 550 nm. White boxes show the images of Fig. 6e.

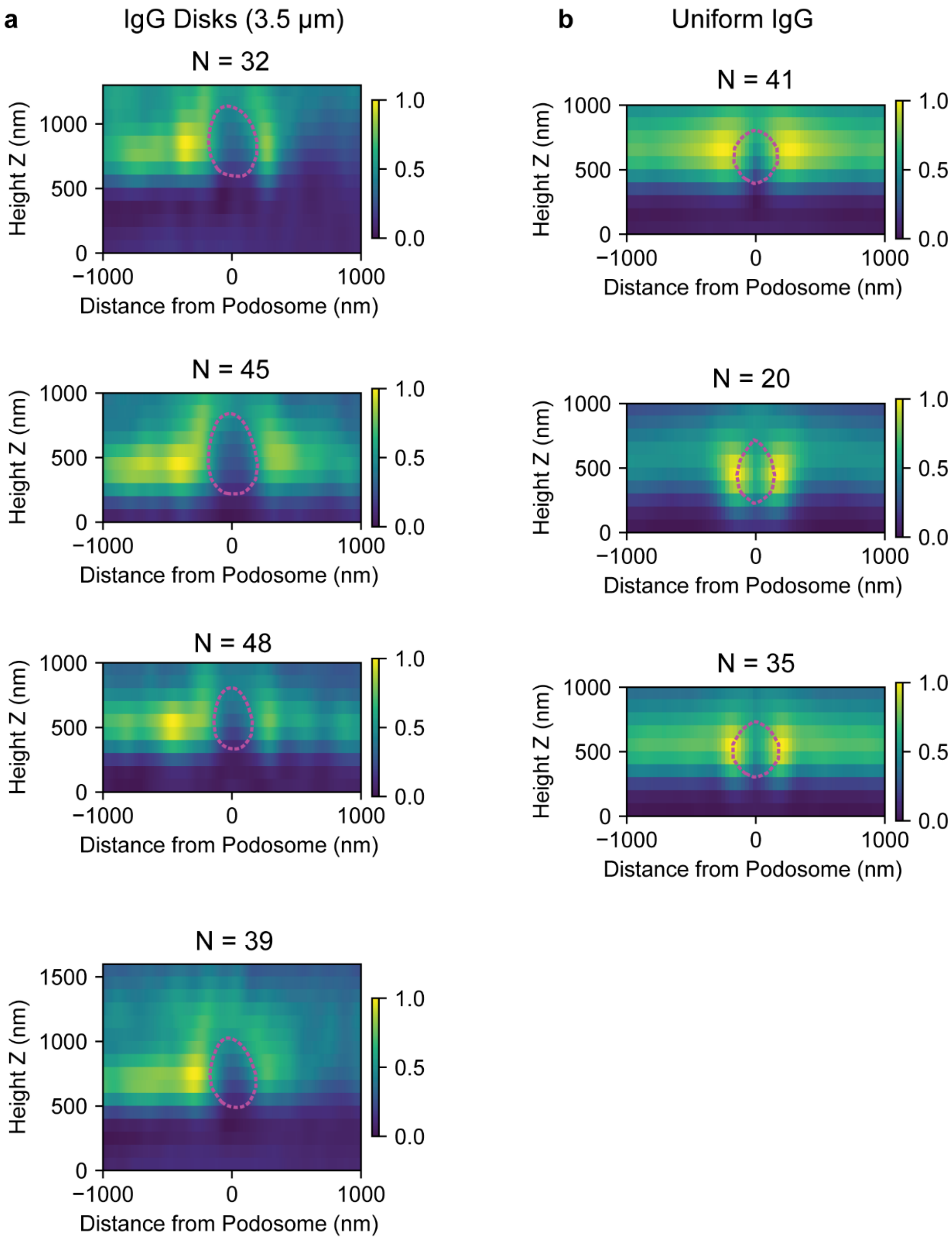


**Supplementary Figure 6. 3D heatmap representations of paxillin. a,b)** 3D-SIM results for paxillin on uniform antibody (**a**) or on IgG disks (**b**). On uniform IgG, the heatmaps are mirrored radial averaging heatmaps, as there is no phagocytosis site center to orient the analysis. On the IgG disks, these are the perpendicular line scan heatmaps. Each heatmap shows a single cell, and N is the number of podosomes. Magenta contour is drawn for the actin channel at 70% maximum intensity.

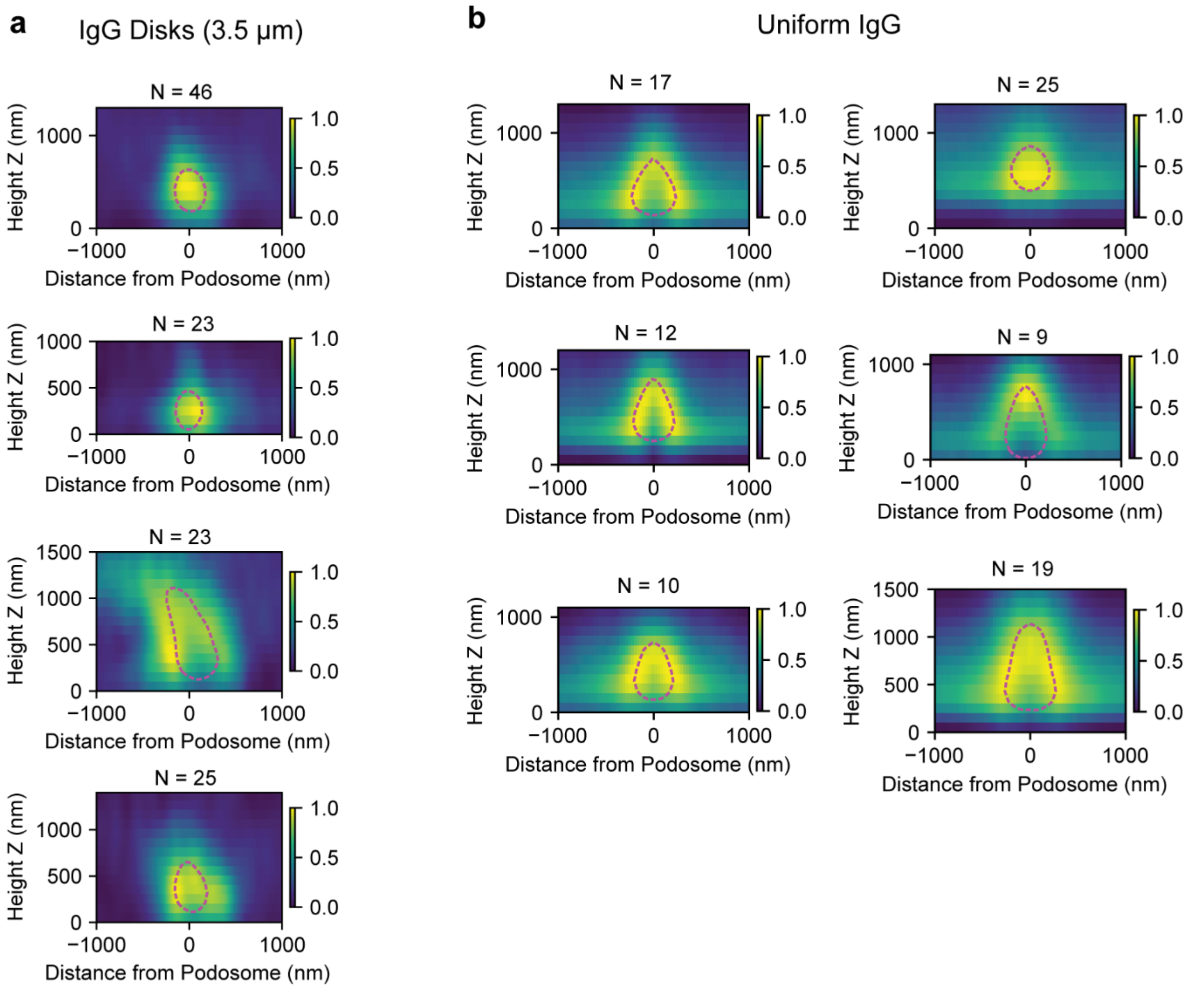


**Supplementary Figure 7. Imaging of α-actinin. a,b)** 3D-SIM results for α-actinin on IgG disks (**a**) or on uniform antibody (**b**). On uniform IgG, the heatmaps are mirrored radial averaging heatmaps as there is no phagocytosis site center to orient the analysis. On the IgG disks, these are the perpendicular line scan heatmaps. Each heatmap shows a single cell, and N is the number of podosomes. Magenta contour is drawn for the actin channel at 70% maximum intensity.

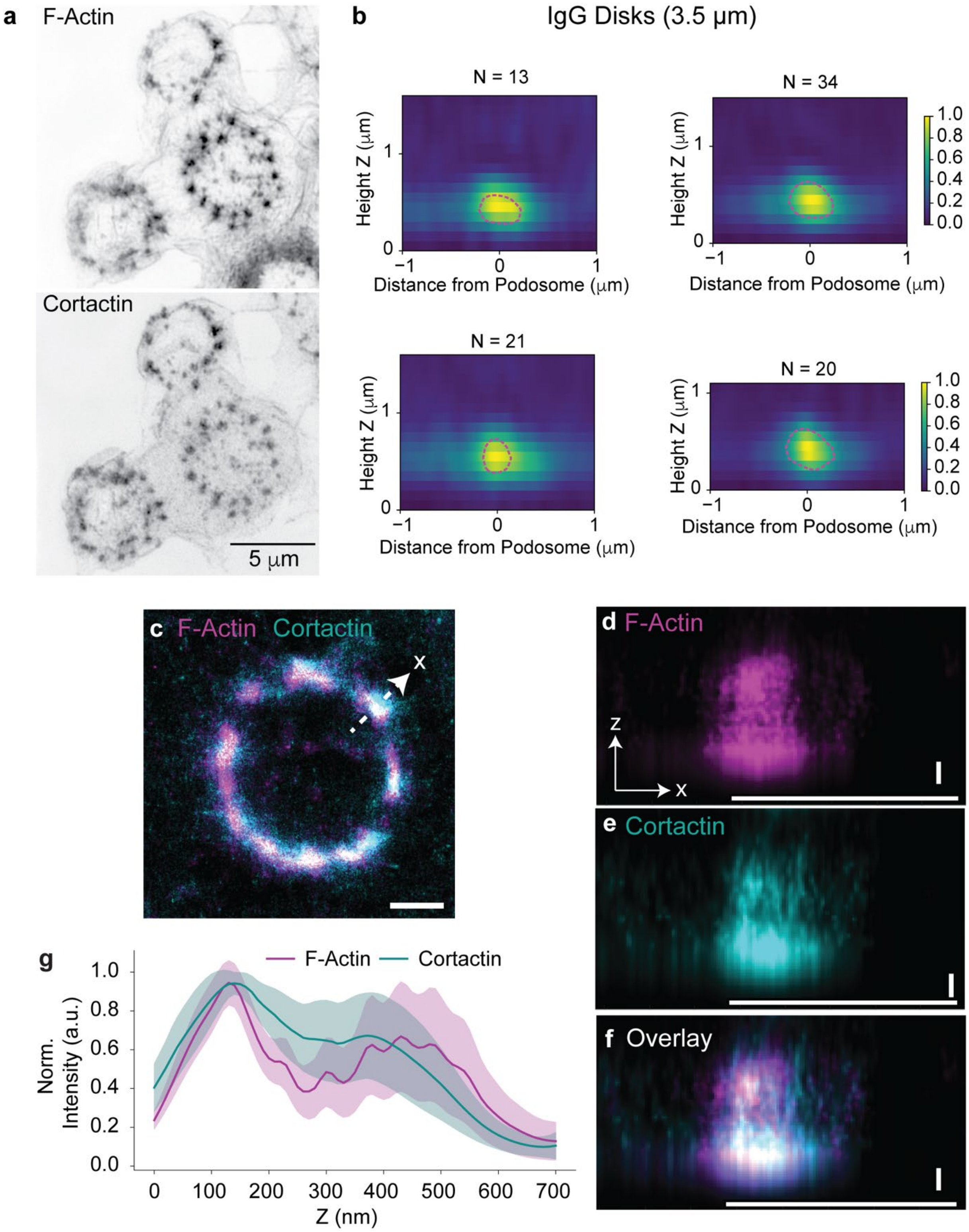


**Supplementary Figure 8. Imaging of cortactin. a)** Z-projections of F-actin (phalloidin Alexa Fluro 568) and cortactin-EGFP from 3D-SIM imaging. **b)** Perpendicular line scan heatmaps of cortactin from 3D-SIM. Each heatmap shows a single cell, and N is the number of podosomes. Magenta contour is drawn for the actin channel at 70% maximum intensity. **c)** Z-projection of F-actin (phalloidin Alexa Fluro 647) and cortactin-mEos3.2 from 3D-PALM/STORM imaging. Scale bar is 1 micron. Representative, manually drawn line scan shown for a single podosome. **d-f)** Heatmaps for F-actin and cortactin distribution for a single podosome, from 3D-PALM/STORM imaging. X-axis scale bar is 1 micron, Z-axis scale bar is 100 nm. **g)** Normalized, mean intensity for F-actin and cortactin from n = 31 podosomes (across 3 cells), from 3D-PALM/STORM imaging.

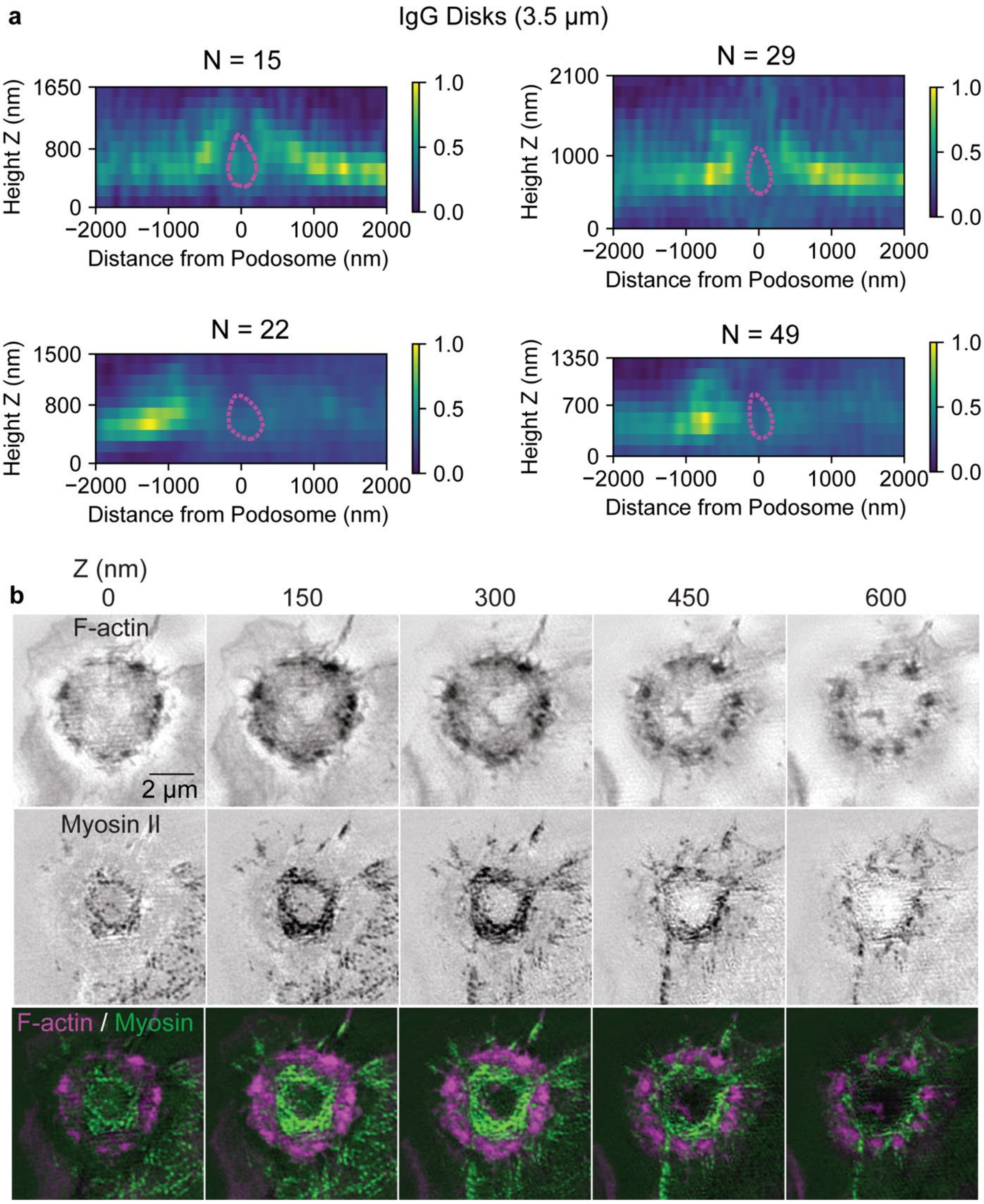


**Supplementary Figure 9. 3D myosin distribution relative to podosomes. a)** 3D-SIM of myosin on IgG disks (perpendicular line scan heatmaps). Each heatmap shows a single cell, and N is the number of podosomes. Magenta contour is drawn for the actin channel at 70% maximum intensity. **b)** Additional 3D-SIM Z-stack images showing actin marked with FTractin-tdTomato, myosin II marked with RLC-EGFP, and merged.

**Supplementary Movies**

**Supplementary Movie 1**. Primary mouse macrophage expressing FcγRIIA-EGFP (green) and Lifeact-mCherry (magenta) migrates over a 15 μm IgG disk, forming podosomes around the edge of the disk. The images were acquired by confocal microscopy at 5 seconds intervals.

**Supplementary Movie 2**. TIRF-SIM of actin podosome ring formation during frustrated phagocytosis. RAW 264.7 macrophages expressing EGFP-FTractin to mark F-actin. The duration of the movie is 4 min and 40 seconds with 5 second intervals between frames using TIRF-SIM.

**Supplementary Movie 3**. iPALM Z stack of a FP site. F-actin is labeled with phalloidin Alexa 647. The colors indicate z distance of actin from 0 (red) to 550 (magenta) nm. There are 10 nm intervals between frames.

**Supplementary Movie 4**. Volumetric visualization of an individual podosome. The visualization is rotated 360 degrees from the bottom to the top along the x axis. There are 2 degrees between frames. Note the actin knob extending from the bottom of the podosome.

**Supplementary Movie 5**. 3D-SIM Z stack of F-actin and myosin II. RAW 264.7 macrophages expressing RLC-EGFP to mark myosin II filaments (green) and FTractin-tdTomato to mark F-actin (magenta). The length of the movie is 1200 nm with 150nm intervals between frames.

**Supplementary Movie 6**. Rotation of a 3D-SIM projection of F-actin and myosin II. RAW 264.7 macrophages expressing RLC-EGFP to mark myosin II filaments (green) and FTractin-tdTomato to mark F-actin (magenta). The 3D projection is rotated 90 degrees from the top to the side along x axis. There are 10 degrees between frames.

**Supplementary Movie 7**. Microtubules inside a circle of podosomes. RAW 264.7 macrophages expressing a GFP fusion of the ensconsin MT-binding domain (EMTB-EGFP) to mark microtubules (cyan) and Lifeact-Halo-549 to mark F-actin (magenta). The duration of this TIRF-SIM movie is 1 min and 18 seconds with 2 second intervals between frames. Note the MT restricted within a circle of actin (approx. 5 o'clock).

**Supplementary Movie 8**. Dynamics of microtubule plus-ends during macrophage frustrated phagocytosis. RAW 264.7 macrophages expressing EB3-EGFP to mark the plus-end of microtubules (green) and Lifeact-Halo-549 to mark F-actin (magenta). The duration of this TIRF-SIM movie is 1 min and 24 seconds with 2 second intervals between frames. This movie shows MT restricted to the interior of a circle of podosomes (approx. 10 o'clock).
